# Articulatory timing and form support distinct neural benefits during audiovisual speech

**DOI:** 10.64898/2026.07.14.738583

**Authors:** Aaron Nidiffer, Aisling O’Sullivan, Edmund C Lalor

**Author notes:** Corresponding Author: Department of Biomedical Engineering, University of Rochester, 201 Robert B. Goergen Hall, P.O. Box 270168, Rochester, NY 14627.

## Abstract

In noisy environments, visible speech articulations improve listening comprehension. The benefit derives from several sources, including articulatory timing and shape. Recent research has shown that visual cortex encodes a categorical representation of articulatory features and that visual speech can benefit both acoustic and phonetic feature processing separately. The present study advances the hypothesis that the shape of the articulators specifically influences the categorization of auditory speech in terms of its phonetic features. We tested this by linearly modeling electroencephalographic responses to natural, continuous speech (in noise) in terms of the acoustic and articulatory features of the speech. We compared the performance of these models in conditions where the speech was accompanied by a natural video of the speaker with their mouth visible, and a video where their mouth was covered by a dynamic ellipse obscuring articulatory shape but preserving dynamics. The dynamic mask reduced comprehension, neural processing of phonetic features, the associated multisensory benefits, and indices of visual-only linguistic processing over occipital scalp. Our findings support substantial visual involvement in speech comprehension, derived largely from the shape of the articulators. They also corroborate several proposals involving audiovisual speech processing hierarchy and the nature of the information contained in visible speech.

**Highlights:**

- Visual speech provides at least two forms of information to enhance acoustic speech processing: redundant temporal dynamics and complementary articulatory information
- Covering the mouth with a dynamic mask preserves horizontal and vertical lip movement information, but largely removes articulatory detail
- Visual speech with a mask preserves some general multisensory benefits but removes visual linguistic information and its ability to enhance auditory processing at the level of phonetic features.

## Background

Listeners understand speech better when they can see the talker producing the utterance. This is especially true in challenging listening environments when speech acoustics are degraded by noise or masked by distracting speech [1,2]. Several properties of audiovisual speech have been highlighted to explain how visual speech improves audio speech comprehension: temporally correlated dynamics of audio and visual speech, visible pre-articulatory movements that are predictive of subsequent articulations, and movements and articulatory shapes that provide complementary information that compensates for what is lost when acoustic speech is masked by noise. These properties have been invoked when explaining audiovisual speech benefits as deriving from various perceptual and cognitive processes such as selective attention [3,4], prediction [5], perceptual filling-in [6], and linguistic constraint [7,8].

Auditory speech processing is accomplished via multiple stages of analysis [9,10] that occur rapidly [11] and in parallel [12] during speech perception. This is often proposed along with another hypothesis positing that each stage of processing feeds back on previous stages, providing the basis for predicting upcoming inputs [13]. Visual speech processing is similarly hierarchical, with evidence suggesting that processing of visual speech spans the same range of linguistic analysis as acoustic speech [14]. Indeed, neurophysiological evidence has supported the presence of both energetic and linguistic representations in the visual system [15–17] and their importance toward processing the information content of speech [18,19, c.f. 20].

The idea of hierarchical speech encoding within both the visual and auditory systems is consistent with the above-mentioned proposals that suggest that different features of visual speech can somewhat independently influences auditory speech processing. Specifically, it has been suggested that 1) the correlated timing of low-level visual speech dynamics provides predictive information about the timing of auditory speech, and 2) higher representations of the shape/form of visual speech provide complementary information about the content of auditory speech. Supporting evidence for these separate modes has been provided – for example – by studies wherein the articulatory details of visual speech stimuli were removed while preserving their dynamics, which resulted in intermediate levels of audiovisual benefits [21,22]. Moreover, this visual-timing benefit is only present if the speech sounds are presented in low levels of background noise [23,24]. These findings suggest that visual timing information serves an important role in audiovisual speech processing but that in higher levels of noise, access to additional complementary visual speech form information is critical.

Recently, our group has begun to identify neurophysiological markers of hierarchical audiovisual speech integration. In one study, we reported evidence that visual speech can independently influence the encoding of low-level acoustic and higher-level phonetic speech features [25]. Specifically, we showed that measures of the encoding of both spectrotemporal and phonetic features were enhanced in audiovisual speech responses relative to what would have been expected from the summation of separate audio and visual speech responses. And we found preliminary evidence that the strength of this multisensory enhancement was more pronounced at the level of phonetic processing for speech in noise relative to speech in quiet, indicating that listeners rely more on articulatory details from visual speech in challenging listening conditions.

In the current study we sought to better understand how such hierarchical audiovisual speech integration might depend on articulatory information in visual speech and its representation within the visual system. Specifically, we tested how removing articulatory shape from visual speech influenced audiovisual speech comprehension and the neural activity related to different levels of linguistic analysis. In doing so we hoped to link several threads of research related to visual speech and audiovisual integration along the speech hierarchy. Our specific approach involved recording electroencephalographic (EEG) signals from participants as they watched natural, connected audiovisual speech where on some of the audiovisual trials participants had access to the full face of the speaker, while on other trials we covered the mouth with a dynamic mask that preserved the timing of the horizonal and vertical mouth dimensions but obscured spatial detail. We anticipated that access to the configuration of the articulators (e.g., the shape of the mouth, the visibility of the teeth, position of the tongue) would activate categorical visual responses and in turn enhance the encoding of phonetic features of that speech. We expected evidence of only modest audiovisual speech integration – driven by temporal correlation – when the mouth was covered by the mask.

## Methods

### Participants

Twenty-six native English speakers (ages mean: 23.6 years, range: 18-40 years; gender: 16 female, 6 male, 4 unreported) took part in the experiment. All participants were free of neurological diseases, had self-reported normal hearing, and had normal or corrected-to-normal vision. Informed consent was obtained from all participants before the experiment, and they received monetary compensation for their time. The study was approved by the Research Subjects Review Board at the University of Rochester. At least some data from four participants was excluded: two participants did not finish the experiment and were excluded from all analyses, one participant did not perform the word detection task and was excluded from behavioral analyses, and, due to technical error, EEG from one subject was corrupted and excluded from neural analyses.

### Apparatus and stimuli

Experiments took place in a dark and sound-attenuated room (IAC Acoustics, North Aurora, IL, US). Visual stimuli were presented on a 26-inch LCD monitor (BENQ XL2411P) operating at a refresh rate of 60 Hz. Audio was presented diotically through Sennheiser HD650 headphones at a comfortable listening level. Stimulus presentation was controlled using Presentation software (Neurobehavioral Systems, Berkeley, CA, US) running on a custom-built PC. Eye position was recorded for the duration of the experiment using the Tobii Pro X3-120 (Tobii AB, Stockholm, SE) eye tracker at 120 Hz sampling rate. The head was stabilized in a chin rest (SR Research, Ottawa, CA). For synchronization, eye tracking data were recorded along with Presentation event time stamps using Lab Streaming Layer (SCCN, San Diego, CA, US).

The speech stimuli were drawn from a collection of videos featuring a trained male speaker. The videos consisted of the speaker’s head centered in the frame with a stationary background. We used Adobe After Effects software to perform detailed face tracking on each of fifteen 1-minute videos. This provided tracking nodes around the perimeter of the face, the eyes, nose, jaw, cheeks, and mouth. The tracking was manually checked to ensure accuracy. The nodes around the face were used to obscure the background with a solid black background, to remove visual contribution from hand movements. Four nodes that tracked the centers of the upper and lower lips and left and right labial commissures (Fig 1a) were used to generate a dynamic ellipse with the commissural nodes as its vertices and upper and lower nodes as its co-vertices. The ellipse formed the aperture of a Gaussian blur and had a black border which resulted in obscuring mouth details from the video (i.e., the teeth, lips and tongue) while retaining horizontal and vertical lip dynamics and overall luminance level and color. (See supplemental materials for example video.) This procedure was used to generate videos for the visual- and audiovisual-masked conditions (Vm and AVm respectively). The intact audiovisual (AV) and visual-only (V, or V-only) conditions had no mask over the mouth. Auditory-only (A or A-only) presentations consisted of a video of a static image of a crosshair at the center of the screen. Note: we will sometimes refer to both visual or both audiovisual conditions together as V(m) or AV(m), respectively.

**Figure 1.**
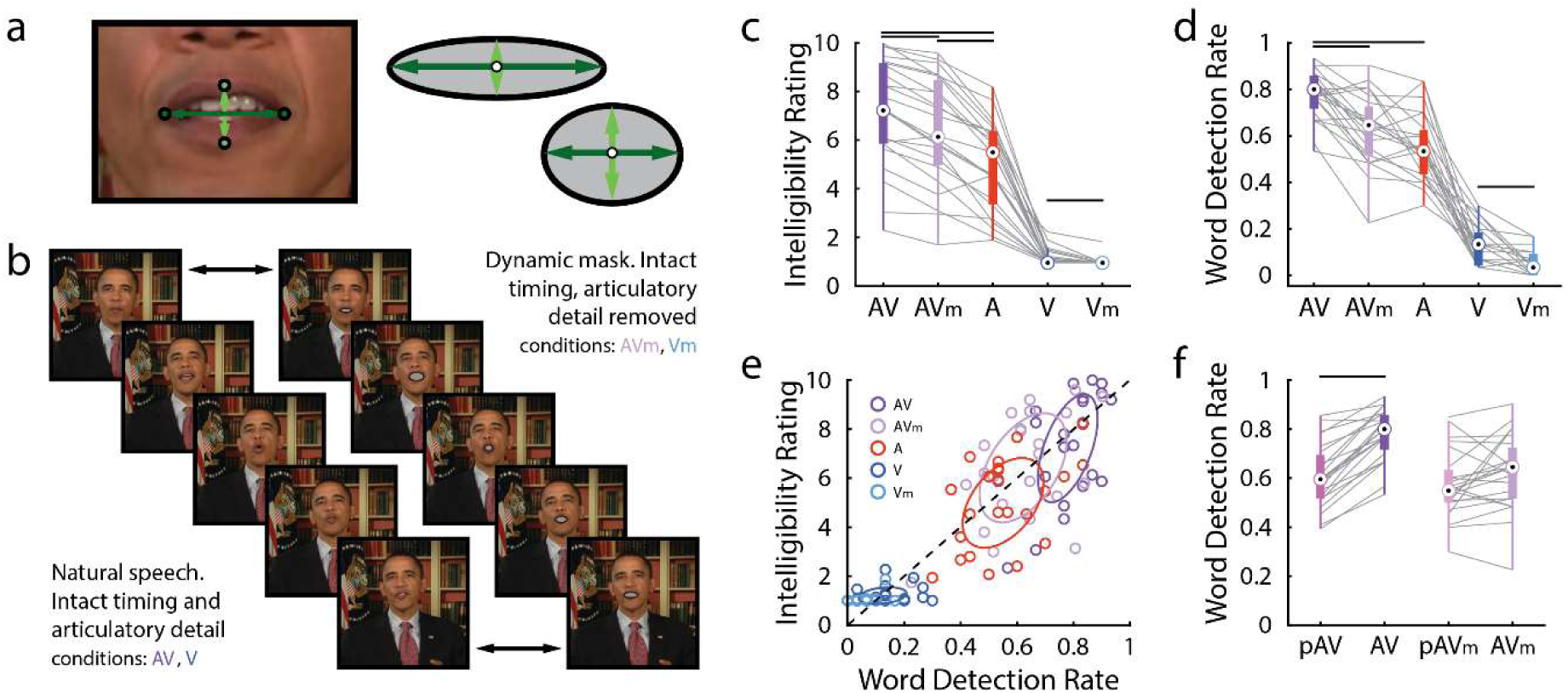
Stimulus manipulation and behavioral results. **a.** We tracked the locations of the lip centers and their lateral commissures on each frame and used these points to construct an elliptical dynamic mask. **b.** The dynamic ellipse followed the lips frame-by-frame, preserving movement timing but obscuring articulatory detail. Intact videos made up visual (V) and audiovisual (AV) conditions while the mask was added to create masked visual (Vm) and audiovisual (AVm) conditions. **c.** Subjective intelligibility ratings for each condition. The dynamic mask consistently gave participants the impression of improved acoustic intelligibility relative to the audio-only condition, while the intact face was judged even more intelligible than the masked face, with and without acoustics present. **d.** An objective measure of comprehension, word detection rate, provided no evidence that the dynamic mask can improve word detection performance, however comprehension improvements from adding articulatory detail remained across audiovisual and visual-only conditions. **e.** Comparison of subjective and objective performance measures. The dashed line represents the line of equality and ellipses are fitted to the horizontal and vertical standard deviations in each condition, with their major axes lying along the first principal component **f.** We measured behavioral multisensory integration by comparing AV performance to a prediction based on probability summation of A and V conditions (pAV and pAVm), which accounts for statistical dependence between modalities. Only when articulatory detail is preserved does AV performance beat this prediction.

All videos then went through the same processing steps. They were first rendered into 1920×1080 pixel movies with a frame rate of 30 fps in Adobe Media Encoder using the Xvid MPEG-4 video encoder (www.xvid.com). The soundtracks from each video were mixed with spectrally matched stationary noise generated using a 50th-order linear predictive model estimated from the original speech recording [26]. Prediction order was calculated based on the sampling rate of the soundtracks [27]. The speech and noise were mixed at an SNR of -7 dB. The speech plus noise audio (or just noise for visual only conditions, with sampling rate of 48 kHz and 16-bit resolution, was then combined with each of the corresponding videos in VirtualDub (www.virtualdub.org).

### Procedure

Participants sat 60 cm from the visual display with their head held stationary in a chin rest as they watched one-minute videos of speech. The experiment was self-paced with participants beginning each video with a button press. During the experiment, each of the 15 speech passages was presented five times, each time as part of one of the five above-mentioned experimental condition: 1) audiovisual speech with visible articulators, AV, 2) audiovisual speech with a dynamic mask over the mouth, AVm, 3) auditory-only speech, A, 4) visual-only speech with articulators visible, V, and 5) visual-only speech with the dynamic mask, Vm, resulting in a total of 75 trials. Presentation order was pseudo-randomized across conditions, ensuring that stimulus repeats (which were presented in different conditions) were spaced at least 15 minutes apart for each participant. No stimulus was repeated within-condition. Participants were allowed free viewing of the stimuli when the visual component was present. On audio-only trials, participants were instructed to fixate the crosshair. They were also instructed to minimize eye blinking and all other motor activity during recording and encouraged to take frequent short breaks between trials.

To encourage active engagement with the content of the speech and to measure speech comprehension, we built in two tasks and analyzed those behavioral data as follows:

1. Prior to each trial, a target word was displayed on the monitor until the participant was ready to continue. The target words would appear 1-3 times per trial, variably to prevent participants from disengaging from the task after they detected the requisite number of targets. Participants were asked to respond to auditory or visual presentations of the target word via button press. A target word was deemed to have been correctly detected (i.e., a hit) if participants responded within 2 seconds after target word onset. We calculated the average hit rate on each condition and for each person. To compare hit rate between unisensory and multisensory conditions, we computed a prediction of multisensory hit rate as the probability summation between the unisensory hit rate while accounting for any spurious decisional enhancements that are due to the possibility that there are two independent channels to infer from [28]. That is, we added their probabilities and subtracted the joint probability:

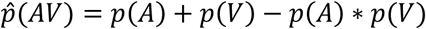

and when the observed multisensory performance exceeds this prediction (*p*(*AV*) > *p^*(*AV*), we interpret this as evidence of multisensory integration.
2. Participants were asked to rate the subjective intelligibility of the speech stimuli at the end of each trial. Intelligibility was rated as a percentage of the total words understood using a 10-point scale (0–10%, 10–20%, … 90–100%). The average rating was computed for each condition and participant.

### Speech representations

Our aim was to examine how vision affects speech processing at different hierarchical levels by measuring separable EEG indices of neural encoding at each level. Following previous work from our lab [25,29] and others [11,30,31], we modeled EEG responses using speech representations derived from multiple levels of the acoustic/visual-linguistic hierarchy. The underlying idea is that EEG responses might reflect the activity of neuronal populations in auditory cortex that are sensitive to spectrotemporal acoustic fluctuations and of neuronal populations that may be invariant to spectrotemporal differences between utterances of the same phoneme or phonetic feature and, instead, are sensitive to the feature category itself. As such, for the present study, we calculated the following representations of the speech signal:

### Envelope and Spectrogram

The acoustic spectrogram was obtained by first filtering the speech stimulus into 16 frequency bands between 80 and 8000 Hz using a compressive gammachirp auditory filter bank that models the auditory periphery [32]. The amplitude envelope for each frequency band was then calculated using the Hilbert transform, resulting in 16 narrow band envelopes forming the spectrogram representation (with dimensions frequency by time). We derived the broadband acoustic envelope (with dimensions one by time) by simply summing each spectrogram across frequency bands.

### Phonetic features

To compute a categorical representation of the speech in terms of its phonetic features, we first identified the start and end of every phoneme in the stimulus. We did this using the Prosodylab-Aligner [33], which, given a speech file and the corresponding textual orthographical transcription, automatically partitions each word into phonemes from the American English International Phonetic Alphabet (IPA) and performs forced alignment [34], returning the starting and ending time points for each phoneme. Manual checking of the alignment was then carried out and any errors were corrected. This information was then converted into a multivariate time-series (dimensions phonemes by time) with a *one* corresponding to the presence of each phoneme at each time point throughout phoneme duration and *zeros* everywhere else. This representation was then converted to describe the articulatory properties of each phoneme (e.g., place, manner, voicing) and generate the main feature of interest for this study, a 19-dimensional phonetic feature representation (feature by time), by mapping each phoneme (e.g., /b/) onto its set of phonetic features (e.g., bilabial, plosive, voiced) as defined previously [35,36].

### Visemes

Visemes were calculated from the phoneme representation. Each phoneme was mapped into phoneme equivalence classes, otherwise known as visemes, which are based on the visual confusability of each phoneme [e.g., /b/, /p/, and /m/ are virtually indistinguishable from visible cues alone; 39]. Depending on the level of skill in lipreading, individuals may be able to discriminate more or fewer visemes. However, we opted to use the 12-viseme mapping defined by [38] for all participants because it balances efficient information extraction and human perceptual sensitivity to visual speech.

### EEG acquisition and preprocessing

Continuous high-density EEG data were recorded using an ActiveTwo system (BioSemi, Amsterdam, NL) from 128 scalp electrodes and two mastoid electrodes at a sampling rate of 512 Hz. Triggers indicating the condition, stimulus number, and start of each trial were recorded along with the EEG. Subsequent preprocessing was conducted off-line in MATLAB. The data were bandpass filtered using second-order, zero phase-shift Butterworth filters between 0.3-30 Hz, downsampled to 64 Hz, and re-referenced to the average of the mastoid channels. Channels contaminated by noise (i.e., with RMS power >3 s.d. above the surrounding channels) were recalculated by spline-interpolating the surrounding clean channels using EEGLAB [39].

### Linear modeling of EEG responses

To relate neural responses to the continuous speech stimuli, we used regularized (ridge) linear modeling to compute a time-resolved stimulus-response mappings – temporal response functions (TRFs) and canonical components analysis (CCA) – that can be used to predict unseen data. We fit TRFs using the mTRFtoolbox (https://github.com/mickcrosse/mTRF-Toolbox), a custom-built toolbox in MATLAB [40,41, but also see 42]. For a more biologically interpretable estimate of this mapping analogous to an event related potential, we fit forward encoding models that predict EEG based on a given stimulus feature. However, encoding models tend to have low sensitivity. For analyses requiring more sensitivity (at some expense of interpretability) we used either a decoding model approach in which we attempt to reconstruct an estimate of the speech envelope from the EEG, or Canonical Components Analysis [43] when we were interested in multi-variable feature mapping. We performed CCA using modified (i.e., adding regularization) functions from NoiseTools toolbox (http://audition.ens.fr/adc/NoiseTools/). We computed the TRF across two sets of time-lags. We used a broader range of time lags – from -300 to 500 ms – for visualization (Fig. 3a), for discriminant analysis (Fig 3b; more details below), and for single lag reconstructions (Fig. 3f,g) in which we attempt to reconstruct a stimulus feature from each lag individually. In line with previous work [19,25,44,45], we used a more restricted range of lags – from 0 to 400ms – for prediction evaluation (e.g., Fig 3a) that integrates information from the EEG across time-lags (and across channels for decoders and CCA). TRF models used for model evaluation (multi- and single-lag) were optimized individually for each subject and condition using leave-one-out cross-validation by 1) fitting the linear model on all trials but one, 2) using the model to predict a new input, 3) evaluating performance on the left-out trial, and 4) rotating the data partition until all trials were left-out for evaluation. Models were evaluated as the Pearson correlation between a prediction and observation across time. We selected the regularization parameter that produced the best prediction accuracy of the left-out data, averaged across all channels and crossvalidation partitions. In the case of CCA, the linear mapping is multivariate and returns a number of correlated components (and their component scores) equal to the smaller of the input and output dimensions (e.g., only 16 components when mapping 128 EEG channels to 16 spectrogram bands). We then take the correlation of the first, most-correlated component as our measure [25]. To prevent any one participant’s data from dominating decoders that were visualized or used in LDA, a common regularization parameter was chosen arbitrarily to be uniformly applied to each participants data for each application (viz. λ=10^5^, LDA λ=10^4.65^).

Under the hypothesis that there is a loss of higher-level visual speech encoding and higher-level audiovisual integration during the masked audiovisual (AVm) condition, we were interested in measuring the differences between A-only responses and AV responses and understand how AVm responses compared to both. Using the MATLAB built-in function *fitcdiscr()*, we used linear discriminant analysis (LDA) to probe whether AVm decoders were more auditory-like or audiovisual-like. Specifically, at each time-lag, we used 80% of auditory and audiovisual trials to train an LDA model to discriminate decoder weights and evaluated its performance on classifying the left-out 20% of A/AV trials. We also trained a null version of this model by shuffling modality labels (A/AV) during the fitting procedure. Using the fitted models, we then classified all AVm trials as either “AV” or “A.” This fitting and testing process was repeated 1000 times, assigning trials randomly into train and test partitions and choosing new random null model labels on each iteration. To visualize model performance, we plot the average predicted class for each time-lag (1 = “AV”, 0 = “A”), or, in other words, the probability that the model classified weights as “AV.” For the out-of-set (AVm) classification, we subtracted the performance of the null model before visualization. To visualize model coefficients (Fig. 3g), we divided the within-class covariance matrix (“Sigma” object property) from the between-class covariance matrix (“BetweenSigma” object property) and extracted the diagonal from the resultant matrix.

For some of our modeling analyses, we wished to separate the effects of one stimulus feature from the EEG before analyzing the effects of another. To this end, following previous work [19,46] we fit a TRF to the to-be-removed feature, predicted EEG using that TRF and the stimulus feature on a given trial, and subtracted the predicted EEG from the observed EEG on that trial. These residuals were then used in subsequent analyses involving other features.

### Indexing multisensory integration

A primary aim of our study was to examine how the available visual information affects participants’ understanding of the speech as well as how it affects our measures of the neural responses to audiovisual speech. Similar to how we look for multisensory effects in behavior (described above), an important aspect of the comparison is to generate an audiovisual baseline that accounts for the separate effects of lipreading (which are present in visual and audiovisual speech) and visual enhancement (which is present in audiovisual speech). That is, we compare the audiovisual responses to the sum of auditory and visual responses [28]. Our five experimental conditions (A, V, AV, Vm, AVm) were chosen with the intention of generating these additive models to compare multisensory effects when articulators are visible – AV≠(A+V) – and multisensory effects when articulators are not visible – AVm≠(A+Vm). For TRF analyses, this involved first obtaining AV(m) TRFs fit on multisensory data (as described above) and A+V(m) TRFs by summing TRFs fit on unisensory A and V data [after 44] and evaluated on their ability to predict AV data. More specifically, the mTRFtoolbox function *mTRFmulticrossval()* fits an A+V(m) model by summing input covariance matrices of A and V(m) EEG and dividing out their average cross-correlation with their respective stimulus features. Like AV, A, and V decoders, A+V(m) models were fit with leave-one-out cross-validation by selecting a single regularization parameter (shared for A and V(m) decoders) that produced the best prediction of AV data, averaged across all channels. For CCA, AV(m) models were obtained by fitting CCA to AV(m) data and A+V(m) models were obtained by fitting CCA to the sum of EEG recorded during A and V(m) presentations of the same stimulus [after 25]. For both TRFs and CCA, A+V(m) models and AV(m) models were then compared on their ability to predict left out AV(m) data. Any performance improvements – AV(m) > A+V(m) – were attributed to nonlinear multisensory effects that assume nothing with respect to their underlying generators. That is, audiovisual speech could affect processing in areas identified by the unisensory decoders, new areas that are only active (or detectably so) in the audiovisual condition, or both.

### Eye Tracking

To ensure that participants were accessing the visible speech articulations, we recorded eye position during each trial. This was especially important to rule out that any differences between masking conditions couldn’t be attributed to gaze. Eye tracking data were analyzed offline. X- and y-position data were down-sampled to 64hz and time-aligned to the video features by segmenting the time-series starting from the trigger sent to Lab Streaming Layer through the duration of the trial, 60 seconds. To account for facial position and size changes across video cuts and head movements, we normalized eye positions into mouth-centered coordinates. Specifically, for each video frame, we subtracted a reference point at the center of the mouth – found by taking the average x- and y-position of the four mouth tracking points – from the instantaneous x- and y-positions and normalized the resulting “mouth-centered” positions by dividing them by the instantaneous horizontal and vertical extent of the oral aperture. A rectangular region of interest (ROI) corresponding to mouth fixation was defined as two times the horizontal and vertical oral aperture on each frame (Fig 2a).

**Figure 2.**
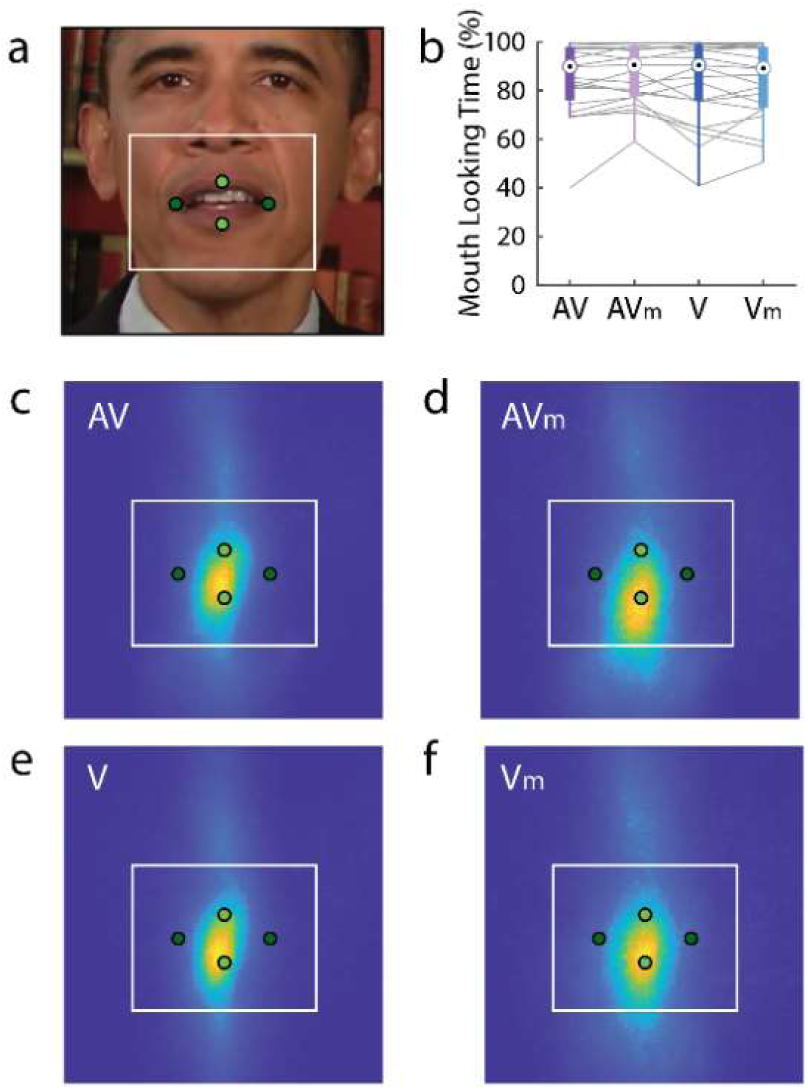
Gaze does not change across visual conditions. **a.** We normalized eye position to the face by subtracting out the center of the mouth tracking points, frame-by-frame. The white box denotes the target region for quantifying mouth looking time – twice the width and height of a box bounding the extremities of the mouth (green dots). These landmarks serve as anchors for panels c-f **b.** Mouth looking time for each visual condition, quantified as percent of time gaze fell within the target box. **c.** Gaze heatmap for intact audiovisual speech, **d.** masked audiovisual speech, **e.** intact visual-only speech, and **f.** masked visual-only speech.

## Results

We analyzed electroencephalography (EEG) from healthy adults as they watched 75 one-minute clips of auditory-only, visual-only, or audiovisual speech in noise (A, V, and AV). To investigate the perceptual and neural effects related to having access to the fine visible detail of the articulators, we added two conditions – visual and audiovisual – in which we imposed an elliptical mask over the mouth area that followed the vertical and horizontal movements of the lips (Vm and AVm; Fig 1a, b), preserving their dynamics. During each trial, participants were looking or listening for a target word which was cued before the start of the trial, reporting detection of the target word via button press. At the end of each trial, they gave a subjective intelligibility rating corresponding to the percentage of words they thought they heard.

### Articulatory timing and form contribute separately speech comprehension

We were first interested in the behavioral consequences of seeing the fine articulatory detail of the speech signal. Previously, it has been shown that visual temporal cues and articulatory detail provide separate benefits to speech perception [21,22]. Unsurprisingly, fully intact audiovisual (AV) speech was judged clearer (Fig 1c; T(22) = 10.28, p = 7.3×10^-10^) and allowed participants to detect target words at a higher rate (Fig1d; T(22) = 10.65, p = 3.8×10^-10^) than acoustic-only (A or A-only) speech. This improvement was also evident when comparing intact AV speech to masked audiovisual (AVm) speech in both subjective clarity (T(22) = 7.57, p = 1.5×10^-7^) and objective detection rate (T(22) = 5.88, p = 6.5×10^-6^), illustrating that the structural detail of the articulators – a feature suggested to carry information about the content of speech [5,22] – confers benefits beyond what their dynamics provide. Importantly, we observed the effect of articulatory detail even in the absence of sound. Participants rated intact visual-only (V) speech clearer than masked visual speech (Vm; T(22) = 2.97, p = 0.007) and were able to visually detect words at a higher rate when they had access to articulatory detail rather than dynamics alone (T(22) = 4.09, p = 4.1×10^-7^). There was conflicting evidence as to whether the dynamic mask provided some benefit to speech perception. Participants rated AVm speech as subjectively clearer than A-only speech (T(22) = 7.09, p = 1.5×10^-7^) but a significant effect on word detection would not survive correction for multiple comparisons (α=0.0125, 4 comparisons; T(22) = 2.64, p = 0.015). It is possible that 23 participants were not enough to detect what might be a subtle effect. But previous work has also shown that the benefits of a purely temporal (non-speech) visual signal only occur within a narrow range of noise levels, which are lower than the level used in the current study [24].

Direct comparisons between AV(m) and A-only behavioral measures do not consider a subject’s ability to detect words based on the visual stimulus alone. As such, directly comparing AV and A-only conditions is limited in its sensitivity to multisensory interactions. To overcome this, we calculated a prediction of AV(m) word detection performance based on a logical OR combination of the respective unisensory performances (i.e., probability summation) that assumes independence of the constituent sensory channels and a winner-take-all decision [28]. Any observed performance exceeding this prediction implies that information in the two sensory channels is combined complementarily. We observed such a non-linear effect for fully intact AV speech (Fig 1f; T(22) = 10.51, p = 4.8×10^-10^), but not when articulatory information was removed (T(22) = 1.85, p = 0.07). Thus, when speech included an intact visual component, visual speech was able to effectively improve comprehension via real multisensory enhancement (i.e., convergence at the perceptual level), compared to the masked condition where improvement could not be attributed to meaningful convergence of information relevant to the task.

### Differences in speech comprehension performance across conditions is not easily explained by gaze

Given the visual differences in the stimuli between our conditions (i.e., no visual stimulus vs masked visual speech vs intact visual speech), we were concerned that participants – who were allowed free viewing of the stimuli when the visual component was present and asked to fixate on the crosshair in auditory only trials – might use different gaze behavior across the conditions and that this might complicate the interpretation of our EEG results, namely that participants weren’t viewing the mouth area on masked trials. To examine this, we measured eye position at each moment in time, normalized and referenced the position to mouth-centered coordinates (see methods for more detail), and calculated the rate at which gaze fell within a region of interest around the mouth area (Fig. 2a white box). We looked at this rate across visual conditions (Fig. 2b-f) and found no evidence for decreased mouth looking time. Specifically, we fit a linear mixed effects model to predict the proportion of time gaze fell within the ROI from the modality (AV vs V), the presence of the oral mask, or their interaction while controlling for stimulus and participant. A significantly large portion of mouth looking time (grand average = 85.4%) could be explained by the intercept (baseline average) term (β = 87.5%, T(1362) = 25.36, p = 1.6×10^-116^), whereas effects for modality (β = 1.8%, T(1362) = 1.44, p = 0.15) and mask (β = 0.9%, T(1362) = 0.69, p = 0.49), or their interaction (β = 1.9%, T(1362) = 1.11, p = 0.27) were small and non-significant. In other words, group performance was largely invariable across conditions. Looking for further support for this, we found that the model performance did not suffer when evaluating reduced models excluding the mask term (LR = 0.48, Δdf = 1, p = 0.49) and when excluding mask and the interaction term (LR = 5.58, Δdf = 2, p = 0.06). The same held true when considering reduced models excluding modality (LR = 2.09, Δdf = 2, p = 0.35), modality and interaction (LR = 2.08, Δdf = 1, p = 0.15), or all three leaving only intercept and nuisance regressors (LR = 6.45, Δdf = 3, p = 0.09). Thus, we conclude that participants were not altering their gaze behavior dependent on either stimulus condition.

### Visual timing and form information separately improves cortical speech tracking

As a first test of how visual speech influences the neural processing of audio speech, we fit decoder models to the EEG signals to evaluate their ability to reconstruct the speech envelope (Fig. 3a). The results largely align with the behavioral findings: Both AV(m) decoder models unsurprisingly outperformed the A model (AV: T(22) = 7.20; p = 3.2×10^-7^; AVm: T(22) = 5.05, p = 4.7×10^-5^) due to the extra visual information present in brain activity, and for both conditions, there was evidence of multisensory integration (i.e., AV(m)>[A+V(m)]; AV: T(22) = 8.18, p = 4.05×10^-8^; AVm: T(22) = 4.03, p = 5.6×10^-4^)), indicating that both an intact face and the ellipse stimulus is being integrated with the speech signal at some level of processing. Finally, envelope tracking was stronger for intact AV speech than AVm speech (T(22) = 3.61, p = 0.0015) suggesting additional activations in response to the added detail of the articulators. This suggestion was supported when inspecting single-lag reconstruction accuracies (Fig. 3 b,c). When comparing AV(m) to A+V(m), cluster-based permutation test identified clusters in both conditions. In the clear visual speech condition, there were two clusters: one occurred just “before” the stimulus (from -200 to -31 ms; T_sum_ = 38.94, p = 0.011) while another larger cluster began at 109 ms and lasted throughout the analyzed time-lags (T_sum_ = 146.60, p = 9.9×10^-4^). In the masked condition, there was one modest cluster spanning 234 to 375ms (T_sum_ = 29.91, p = 0.023).

**Figure 3.**
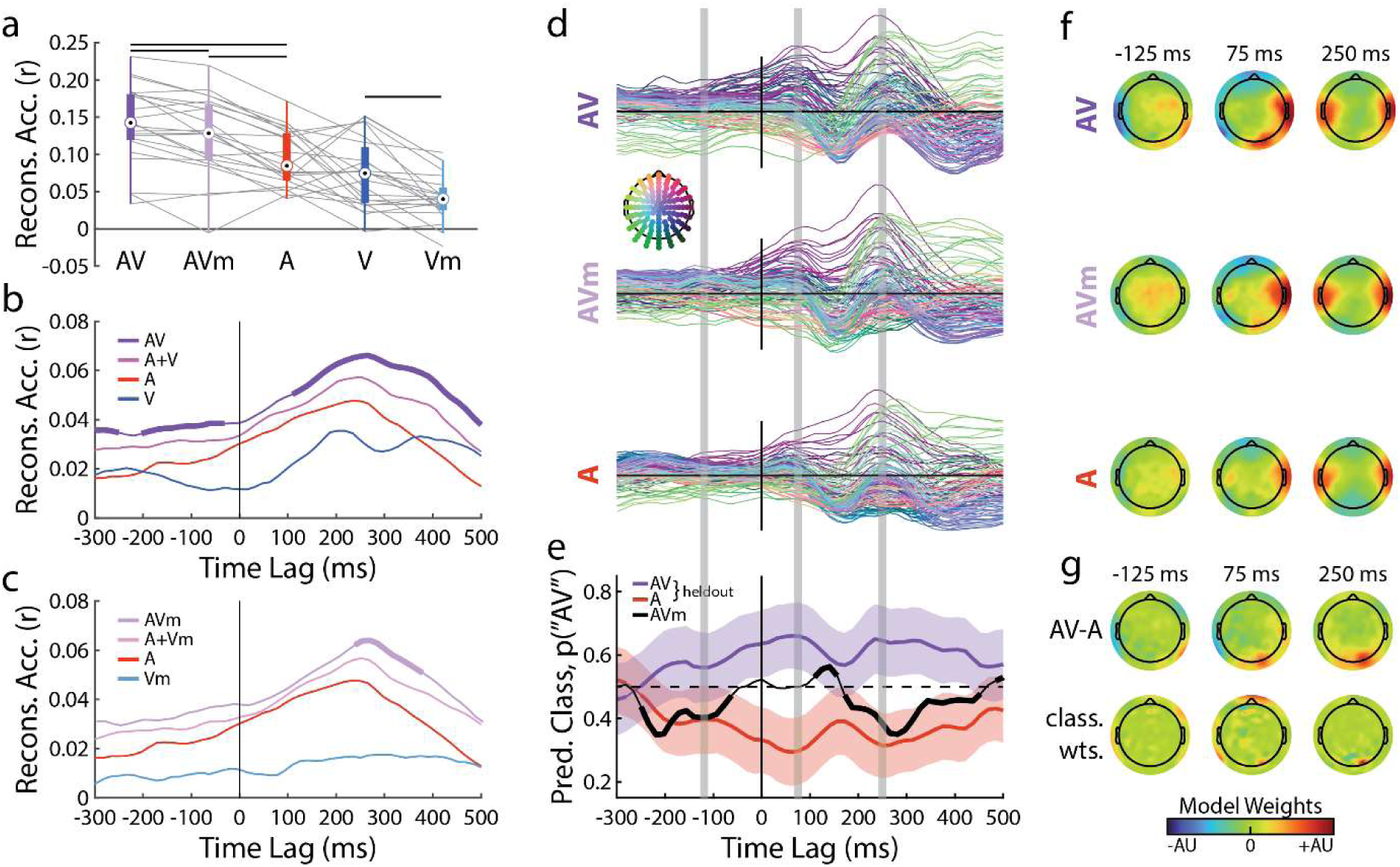
Envelope tracking. **a.** Envelope decoder model weights for conditions containing acoustic speech. Channels are color coded according to the scalp map inset. Grey bars indicate timepoints for inspected topos in c and d. **b.** Decoder classification. We trained a classifier to discriminate AV from A decoder weights at each time lag. We then tested AVm decoders and held-out AV and A decoders. Thick line segments indicate significant clusters that differ from chance performance. **c.** Scalp maps for each decoder and at teach time point shown in panel a. **d.** Decoder difference topographies (AV-A; top) and classifier weights (bottom) for classification shown in panel b. **e.** Envelope reconstruction accuracy for each condition and lags 0 – 400 ms. **f.** Single-lag reconstruction accuracy for auditory and intact visual conditions. Thick line segments indicate significant clusters of time-lags comparing AV vs A+V. **g.** Same as panel f, for masked visual conditions.

To get a deeper insight into the spatiotemporal patterns driving envelope reconstruction, we visualize the decoder weights for A, AVm, and AV speech, resolved over time-lag (Fig 3d) and spatially over the scalp at selected time-points (Fig 3f). The A and AV decoder models appear qualitatively different across channels and time, with the AVm model being intermediate. To quantify the differences between the decoder models – and to test how similar the AVm model was to the A and AV models, we implemented a classification analysis. Specifically, at each time-lag, we applied LDA (Fig 3e) to discriminate A from AV and used cluster-based statistics [47] to test for clusters across time-lags. The classifier was able to reliably discriminate the effects of visual speech on unseen A and AV decoders starting from -250ms “before” speech, likely related to visible articulations that precede the onset of speech sounds [48], through the entirety of tested time-lags (T_sum_ = 3999.6, p = 9.9×10^-4^). We then asked how imposing the dynamic mask over the mouth affected the decoder models. We used the abovementioned discriminant models to classify AVm decoders and found three clusters of time-lags where classification performance was different than performance of a null discriminant model (Cluster 1: -250 to -62 ms; Cluster 2: 109 to 156 ms; and Cluster 3: 172 to 453 ms). Before speech onset, AVm response functions were judged to be auditory-like (T_sum_ = 217.26, p = 9.9×10^-4^). The classifier was then unable to reliably label the earliest (p50-like) cortical responses better than chance. A brief cluster around a putative N100 response was modestly audiovisual-like (T_sum_ = 36.37, p = 9.9×10^-4^). Finally, the latest and largest cluster more closely resembled auditory responses (T_sum_ = 287.06, p = 9.9×10^-4^). The TRF scalp distributions of the effect of visual speech (AV-A) and the classifier weights unsurprisingly have peaks over occipital scalp, becoming stronger for later time-lags (Fig. 3g). Taken together, these results suggest that the ellipse was properly integrated with the speech signal and that visual dynamics and form information may be separately integrated along the speech hierarchy. However, basing this conclusion on envelope tracking is rather indirect. To more directly explore this issue , we next sought to examine how the EEG data reflected the processing of different hierarchically organized speech features.

### Visual form information improves audiovisual processing of phonetic features

Behaviorally, visual speech timing and form seem to affect speech and non-speech items differently in a lexical decision task [22] – visual timing information can improve response accuracy and speed at the identification of speech sounds and non-speech sounds (spectrally rotated speech), while visual form information has an added benefit only for speech signals. This highlights the general property of timing information and implies that form provides some linguistic information that aids in speech recognition. Based on this idea, we hypothesized that visual speech dynamics would have effects restricted to low-level acoustic or temporal processing and the form of the articulators would improve phonetic feature encoding. To have a better understanding of these underlying acoustic and linguistic processes at play during audiovisual speech integration, we turned to multivariate decoding methods by which we can relate multivariate features (spectrogram bands or phonetic features) to multivariate EEG. We performed Canonical Component Analysis [CCA; 43] to quantify multisensory EEG responses related to the acoustic spectrogram and phonetic features (Fig 4). This analysis involved decoding acoustic and linguistic speech features from the full AV EEG. We fit these CCA models to AV(m) EEG and A+V(m) EEG and tested them on left out AV EEG to determine the amount of multisensory integration for each speech feature. Expectedly and consistent with our envelope tracking results, we find multisensory effects persist across visual conditions and feature (Fig 4a, b; AV vs A+V, sgram: T(22) = 6.94, p = 5.69×10^-7^, ph. fea.: T(22) = 4.76, p = 9.44×10^-5^; AVm vs A+Vm, sgram: T(22) = 9.2, p = 4.83×10^-9^, ph. fea.: T(22) = 7.04 p = 4.57×10^-7^).

**Figure 4.**
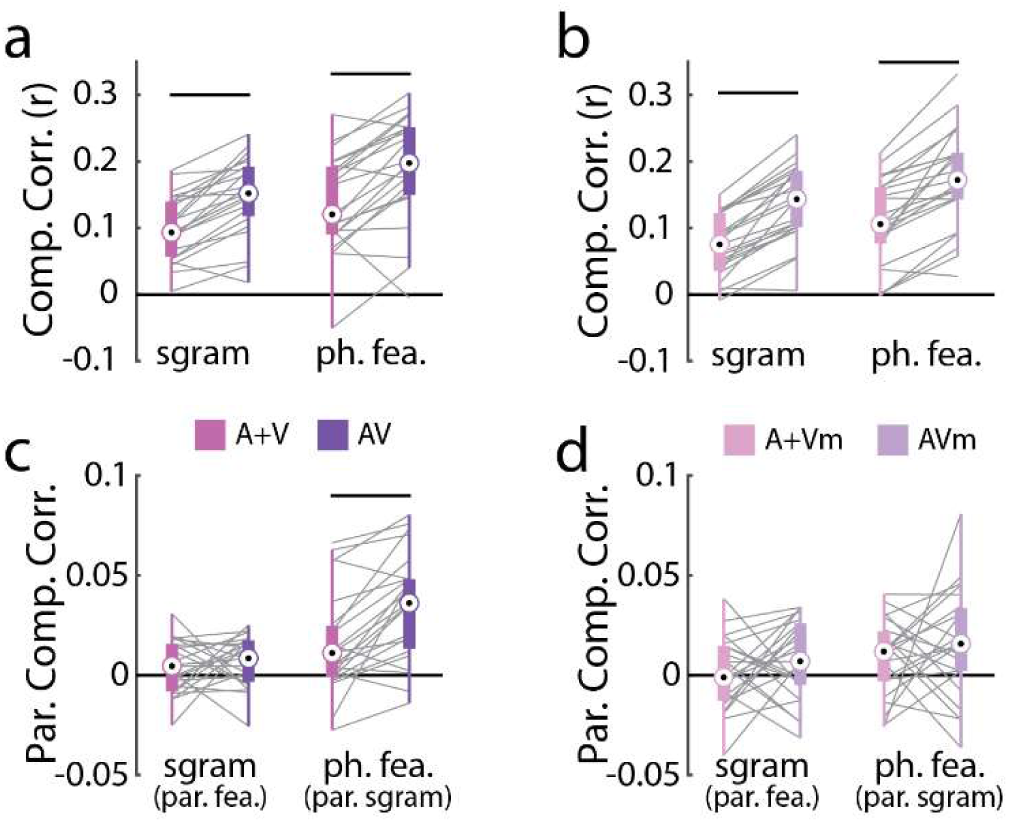
Canonical Components Analysis. **a.** Component correlations for AV and A+V EEG rotations with spectrogram and phonetic feature models, using the full EEG signal. **b.** Same as panel (a.) but AVm and A+Vm EEG **c.** and **d.** Same as panels a and b, but after partialing out the other feature of non-interest.

An important caveat when modeling how EEG responses relate to different speech representations is that those representations are often strongly correlated with each other and may share some explanatory capacity of the EEG – hence the ambiguity of the envelope tracking results discussed above. This makes it difficult to distinguish between, for example, EEG activity that is related to the processing of speech acoustics from EEG activity that is related to the processing of phonetic features. To account for the explanatory power of a given feature that might be shared with a feature of interest, we used an encoding model approach to selectively remove activity related to the feature of non-interest before performing the above CCA analysis on the residual EEG. In the case of masked speech, this approach produced no evidence of multisensory integration (AV(m) > A+V(m)) for the spectrogram after removing EEG activity related to the phonetic features (Fig. 4d, left; T(22) = 1.12, p = 0.27) or for the phonetic features after removing EEG activity related to the spectrogram (Fig. 4d, right; T(22) = 0.95, p = 0.35). However, for intact speech, while there was no evidence of multisensory integration for the spectrogram (Fig. 4c, left; T(22) = 0.49, p = 0.62), there was a significant index of multisensory integration for the phonetic features that remained after removing the EEG activity related to the spectrogram (Fig. 4c, right; T(22) = 4.67, p = 1.15×10^-4^). This suggests that seeing the momentary configuration of the speech articulators enables a listener to encode the categorical phonetic features of audio speech whereas simple access to their dynamics does not.

### Articulatory detail enables the encoding of visemes in visual cortex

In previous work, we identified visemes as a representation of visual speech that is tracked in visual cortex in a way that relates to visual word recognition [19]. Based on this work and current findings suggesting much of late multisensory effects could be attributed to visual rather than auditory generators (i.e., Fig. 3e,g), in the present exploratory analyses we have hypothesized that the encoding of phonetic content in visual cortex is dependent on seeing the shape of the articulators and the physical interaction between tongue, teeth, and lips. After first partialing out activity specific to the timing of the speech stimulus (i.e., that related to the speech envelope), we used encoding models fit to the spectrogram and visemes to predict EEG responses across the scalp (Fig. 5a,c) and focused our analysis on a subset of electrodes over visual cortex (Fig. 5, legend inset). As expected, in both visual and audiovisual conditions, visemes could predict EEG, while controlling for stimulus timing, more strongly when the lips weren’t masked (Fig. 5b,d right panel; AV vs AVm: T(22) = 3.58, p = 0.0017, V vs Vm: T(22) = 3.82, p = 9.31×10^-4^), supporting the hypothesized link between articulatory detail and visual phonetic information. As a comparison, we performed the same analysis using the spectrogram feature. In this case, the model performance did not differ enough between masked and intact audiovisual speech to reach significance (Fig. 5b left panel; T(22) = 1.85, p = 0.07) while in the visual conditions it did (Fig. 5d left panel; T(22) = 3.18, p = 0.0043). Our *post-hoc* interpretation of this finding is that frequency specific information is still (weakly) represented over occipital scalp on AVm trials due to either communication between auditory and visual cortices [49], visual correlates that give a sense of speech’s spectral content [6], or a more inert reason: volume conduction of auditory responses, although if that’s the case we ought to see the same pattern for visemes.

**Figure 5.**
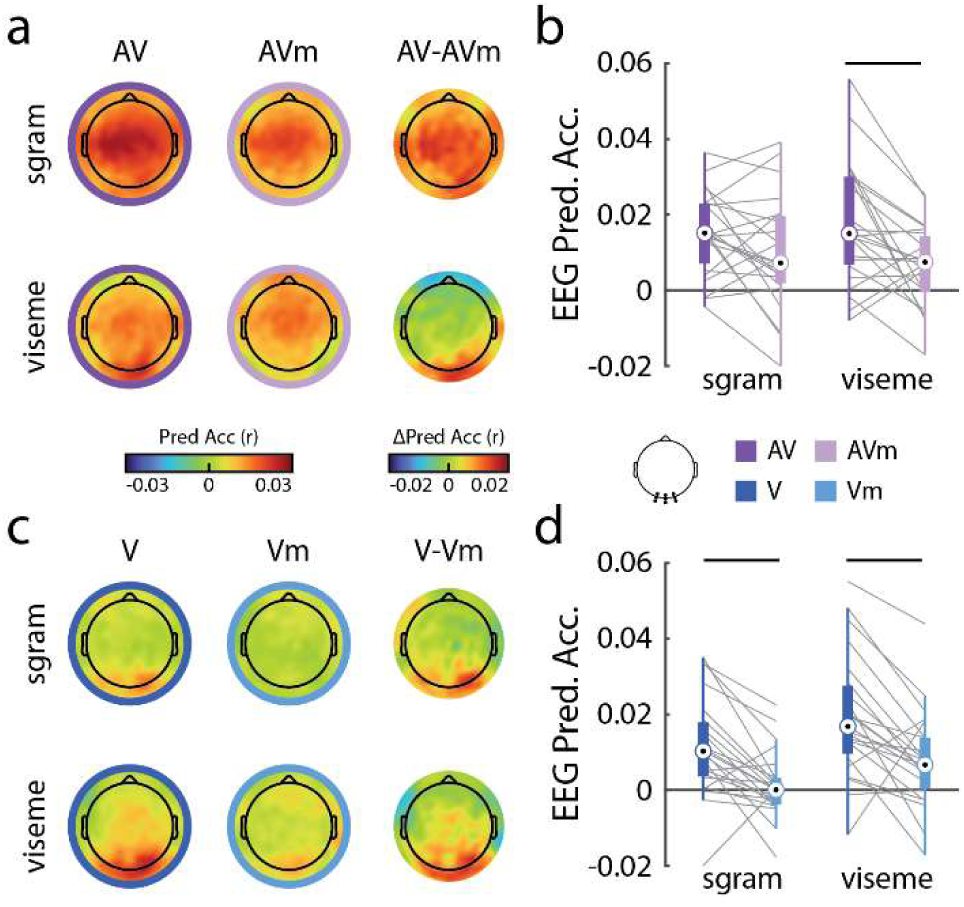
Viseme encoding over occipital scalp. **a.** Scalp distributions of EEG prediction accuracy of the spectrogram feature model and viseme feature model in clear and masked audiovisual speech, and their differences. **b.** Average prediction accuracy for electrodes over visual scalp (see legend inset) for the data in panel (a). **c.** and **d.** Same as panels (a) and (b), but for clear and masked visual speech.

## Discussion

In the present study, we investigated how visible lip dynamics and articulatory detail contribute to speech comprehension and the neural responses to visual and audiovisual speech in noise. Previously work has shown 1) that visual speech dynamics and form have separable effects on speech processing [22]; 2) that, much like acoustic speech, visible speech signals are processed at distinct hierarchical levels [15,16] with comprehension being more tightly linked to higher-level linguistic processing [19]; and 3) that visual speech impacts speech processing seemingly distinctly across multiple levels [25]. In the current study we have woven these threads together by showing that visual phonetic responses are driven largely by the form of the articulators, as previously hypothesized, and that these responses are the major contributors to audiovisual integration at the phonetic feature level leading to improvements in comprehension. We have also demonstrated that integration of common temporal information is present regardless of the presence or absence of articulatory detail. Overall, our results are consistent with the ideas that the visual speech system is organized hierarchically in a way that derives predominantly from distinct features of visible speech articulations and influences the processing of audio speech along multiple, corresponding levels of the linguistic hierarchy.

### A multitude of audiovisual interactions during speech perception

Our findings support a multi-faceted and hierarchical audiovisual speech system. The presence or absence of visible articulatory detail did not appear to provide any unique multisensory benefits at level of acoustic process. However, this manipulation had a profound impact on multisensory integration at the level of phonetic feature processing. This dissociation was likely pronounced, by design, due to our speech stimuli being presented in a high level of background noise [25]. Indeed, with few specific exception [e.g., 6], the dynamics of the lips seem to have little potential for enhancing speech comprehension in moderate to high levels of noise (also see below). Instead, the spatial processing of vision is sensitive to subtle articulatory details that the auditory system struggles with [e.g., place of articulation, 50]. According to our results, access to these visual form cues gives rise to viseme responses in visual cortex and in-turn supplements and/or constrains auditory phonetic feature responses, meaningfully toward improved comprehension in the listener.

The idea presented here of a multitude of audiovisual interactions aligns well with previous neural findings. We and other groups have shown that the influence of visual speech varies across linguistic level, goal, neural mechanism, visual feature, and time-scale [8,25,51–53]. This work also aligns well with a strong base of behavioral experiments. One relatively consistent finding is that altering spatial detail from visual speech results in appreciably lower levels of comprehension benefits [21,22,54–56], a finding we replicate. What is interesting is that, like our findings, the literature is equivocal on gains that can be attributed to visual temporal information – with some studies finding only modest comprehension gains from stimuli involving no face [21,23,24] and others very convincingly unable to find an effect [54,55]. Critically, in these studies, comprehension benefits are observable only in low noise levels (−3 or -1 dB SNR) or provide only a modest (<1 dB) but reliable boost to the speech reception threshold. The abovementioned studies that were unable to find a visual effect were conducted in noise at -8 and -12 dB while the present study was conducted at -7 dB SNR, which could explain the discrepancy. A considerable amount of effort has gone into understanding the role of correlated audiovisual dynamics in multisensory perception [57–60]. Though the effects were largely tested using non-speech stimuli, which alludes to the general nature of the potentially associated cognitive process, they appear consistent between speech and non-speech stimuli [22,61,62] and may reflect a general multisensory computation [63]. Temporal correlation does support multisensory binding and object formation [64] and thus comprehension benefits from lip dynamics may be largely derived from object-based attentional enhancements [c.f., 6]. This evokes two important points: first, it could explain why effects are restricted to low noise levels (but not observed in clear speech) – where visual enhancement of auditory processing is possible (i.e., auditory processing is not at “ceiling”), but the audio isn’t also too degraded to benefit from that visual influence; and second, if the effects are largely attentional perhaps the effects of visual timing cues will be less apparent under some experimental conditions (e.g., highly degraded or low SNR single speaker experiments) compared to others (e.g., distracting speakers/cocktail party paradigms). More work is needed to uncover these answers. Such future work might also consider including other sources of information from vision such as visual head start [5,65] and the availability of linguistic information outside the mouth area [66–69]. It might also benefit from using neurophysiological recording methods that enable exploration of the (possibly separate) anatomical pathways that allow visual speech to influence auditory speech processing [69].

### Complementary visual speech constrains phonetic inference

The above ideas are largely consistent with three proposals regarding vision’s effects on speech processing. In one proposal [70], Campbell divides visual speech cues into two types: *correlated features* that provide redundant information to auditory signals as a backup, and *complementary features* that contain information that, in the auditory modality, is not well resolved in the presence of noise (e.g., place of articulation). Separately, Peele and Sommers propose that vision provides *prediction* of upcoming speech articulations to the auditory system, and *constraint* regarding the linguistic inferences that are drawn from degraded speech. It’s intuitive to see how signals that are correlated have the ability to predict one another when one happens first, as vision mostly does [48], and how two complementary (i.e., non-overlapping) signals have the capacity for constraint [7]. Finally, Kim and Davis [22] frame the visible portions of speech in terms of its *timing* or its cadence, and its *form* of the articulators, or the spatial configuration and interaction of the tongue, teeth, and lips. The analysis framework and resulting findings of the present study are largely consistent with all three of these proposals. We show that temporal correlations from the masked speech provide only modest behavioral benefits and only modest neurophysiological signatures of multisensory integration. Meanwhile, we relate articulatory structure to the encoding of visemes, which implicitly captures the complementary nature of visual speech, and also relate it to multisensory enhancement of phonetic feature responses, which we contend is a consequence of linguistic constraint imposed by vision.

Our results speak to the importance of fine spatial detail in visual speech and the complementary information that it provides in challenging listening conditions. Although implicitly we have coarsely dichotomized the visual speech concepts of timing/form and redundant/complementary along a common divider, not all visual dynamics are redundant with the acoustic dynamics. Although we have had success modeling visual speech using the acoustic envelope here and previously [15,44], lip dynamics provide slightly different temporal information [20]. In addition, using a low-dimensional representation of speech dynamics – such as the speech envelope – to analyze speech responses will often conflate several levels of the linguistic hierarchy that are simply correlated with that input feature [71]. This can lead to misunderstanding about the nature of “cortical speech tracking” and its relationship to comprehension when, in reality, the relationship between comprehension and envelope tracking all but disappears when one accounts for features representing abstract linguistic categories [29–31]. Our prior work [19] and the current results suggest vision and visual speech work analogously to audio speech and should be treated with the same nuance [c.f., 20]. Finally, visual articulatory detail – especially its faithful representation in the stimulus – becomes crucial amidst recent efforts to develop visual avatars for their potential to improve speech comprehension, especially as a form of hearing aid in the hard of hearing [72–78].

## Acknowledgements

This work was supported by NIH grant DC016297 to ECL. Additional support was provided by a Science Foundation Ireland Career Development Award (CDA/15/3316) to ECL. We are thankful to Drs. Jin Dou and David Hernández-Gutiérrez for helpful conversations and feedback.

## Author Contributions

ARN: Conceptualization, investigation, formal analysis, writing – original draft, writing – review and editing, visualization. AOS: Conceptualization, investigation, formal analysis, writing – original draft. ECL: Conceptualization, writing – review and editing, supervision, project administration, funding acquisition.

## Conflict of Interests

The authors declare that they have no conflict of interests, financial or otherwise.

## Notes

### Competing Interest Statement

The authors have declared no competing interest.

